# Stepwise Bayesian Machine Learning Uncovers a Novel Gene Regulatory Network Component in Neural Tube Development

**DOI:** 10.1101/2024.08.25.609396

**Authors:** Chen Xing, Yuichi Sakumura, Toshiya Kokaji, Katsuyuki Kunida, Noriaki Sasai

## Abstract

Recent advancements in machine learning-based data processing techniques have facilitated the inference of gene regulatory interactions and the identification of key genes from multidimensional gene expression data. In this study, we applied a stepwise Bayesian framework to uncover a novel regulatory component involved in differentiation of specific neural and neuronal cells. We treated naive neural precursor cells with Sonic Hedgehog (Shh) at various concentrations and time points, generating comprehensive whole-genome sequencing data that captured dynamic gene expression profiles during differentiation. The genes were categorized into 224 subsets based on their expression profiles, and the relationships between these subsets were extrapolated. To accurately predict gene regulation among subsets, known networks were used as a core model and subsets to be added were tested stepwise. This approach led to the identification of a novel component involved in neural tube patterning within gene regulatory networks (GRNs), which was experimentally validated. Our study highlights the effectiveness of in silico modeling for extrapolating GRNs during neural development.

## Introduction

The neural tube is the embryonic organ of the central nervous system, and is comprised of a number of different types of neural progenitors and neurons arrayed in an orderly manner (Sagner and Briscoe, 2019).

In the pattern formation of the neural tube, the dorsal-ventral (D-V) arrangement is achieved by the gradients of the morphogens emanated from the roof plate and floor plate, which are located at the dorsal- and ventral-most domains of the neural tube, respectively (Le Dreau and Marti, 2012). BMP and Wnt signals emanate from the roof plate, while Sonic Hedgehog (Shh) from the floor plate, and each of them forms a concentration gradient through the D-V axis (Sagner and Briscoe, 2019). Each neural progenitor cell in the neural tube receives these morphogenetic factors, recognizes the concentration of signaling molecules as positional information, and acquires a specific cell fate.

The neural tube is divided into more than ten distinct domains along the D-V axis, each containing specific types of neurons with unique functions (Alaynick et al., 2011; Ribes and Briscoe, 2009). Each neuronal domain corresponds to its progenitor domain, whose positional identity is defined by the concentration gradients of signaling molecules such as BMP, Wnt, and Shh. These domains are characterized by the transcription factors specifically expressed within them. For example, PAX7 and PAX6 are expressed in the dorsal regions where Shh is absent or present at low concentrations. In contrast, genes like NKX6.1, OLIG2, NKX2-2, and FoxA2, which are induced by Shh and contain Gli-binding sequences in their regulatory regions, are expressed in the ventral domains. The broad expression of NKX6-1 in ventral domains suggests that its expression can be triggered by low levels of Shh, while OLIG2, NKX2-2, and FoxA2 require higher concentrations of Shh for their expression (Ribes et al., 2010). The differential expression of these genes indicates that their induction is dependent on the concentration of Shh. Moreover, these transcription factors (TFs) engage in mutual regulation, often through repressive interactions (Briscoe et al., 2000). For instance, OLIG2 and NKX2-2 repress each other’s expression, and both are negatively regulated by PAX6, contributing to the specification of identities along the D-V axis (Balaskas et al., 2012; Dessaud et al., 2010). NKX2-2 also represses PAX6 and IRX3, thereby reinforcing ventral neural identities (Sagner et al., 2018). These interrelationships are collectively referred to as gene regulatory networks (GRNs), and individual GRNs have been discovered through significant efforts by using traditional experimental methods such as overexpression analysis, genetically mutant mice, and biochemical techniques (Balaskas et al., 2012). Nevertheless, given the large number of domains in the neural tube, there should be many more unknown GRNs (Delile et al., 2019). Recently established mathematical methods are expected to play an important role in the comprehensive prediction of GRNs.

A number of methods have been proposed to clarify the correlations between multidimensional gene expression data; however, a definitive method has not yet been established because of the noise often present in the data (Badia et al., 2023; Chen and Mar, 2018; Pratapa et al., 2020). When the number of expression data samples is large, the machine learning-based method GENIE3 shows relatively good estimation performance (Huynh-Thu et al., 2010). On the other hand, when the number of samples is small, iRafNet, which estimates GRNs using metadata such as protein interactions and transcription factor-DNA binding, is superior (Petralia et al., 2015). In addition, the Bayesian approach algorithm improves performance by introducing known gene and pathway information as prior information and constraining the estimation of GRNs (Gao and Wang, 2011; Liu et al., 2016; Wu et al., 2019). These results show that the use of known information is effective under conditions with a small number of samples. In the case of the neural tube formation, at least 10 regulatory genes are known, and some of their pathways have also been elucidated. However, conventional methods, including GENIE3 and the Bayesian approach, analyze data from all genes at once; therefore, there is a possibility that the analysis results may become unstable due to noise or overfitting to a group of genes with low relevance. When there is some known information, such as the GRN in the neural specification, it may be effective to set the known genes and pathways as the core network, and evaluate the validity by adding one of the remaining many genes stepwise. In machine learning, there is a method called the forward wrapper, which tests whether adding a single factor improves performance (Kohavi, 1997).

In this study, we conducted bulk mRNA sequencing of neural progenitor cells cultured at varying concentrations and durations, and employed mathematical analysis to infer unknown genes and pathways. The neural progenitor cells were prepared as intermediate neural explants from the neuroepithelium of the stem zone in the closing neural plate, which are considered to be in a naive state (Delfino-Machin et al., 2005). These cells allow for the direct observation of cellular responses, such as gene expression, to extracellular stimuli like signaling molecules or chemical compounds (Dessaud et al., 2008; Sasai et al., 2014).

Next, to treat similar genes as a group, we clustered the measured gene expression time series for each of the four Shh concentrations, created subsets for genes that belonged to the same class at each of the four concentrations, and analysed the average time series within these subsets. We developed a novel method called “network assembly and inference using stepwise testing with optimization (NAISTO)”, which is a Bayesian statistical learning method that is an advanced form of the elastic net method using known genes and pathways as a core network, replaces and evaluates candidate genes one by one, and assembles a network of necessary genes. When we evaluated the prediction performance of NAISTO using 9 out of 10 known genes as training genes and 1 as a test gene, all 10 test genes were predicted in the top positions. Finally, we predicted the 11th unknown gene from the 10 known genes, and found that one of the top-ranked genes has the activity to change neural tube patterning upon forced expression. These results indicate that the stepwise assembly of GRNs from known genes and pathways is effective to predict new GRNs during the neural tube development.

## Results

### Neural explants cultured with different concentrations and durations of Shh separate the neural and neuronal identities

To systematically replicate the diversity of cell types and differentiation stages induced by Sonic Hedgehog (Shh), we utilized neural explants composed of naive neural progenitors, which serve as an effective system to correlate extracellular inputs with cellular outcomes (Dessaud et al., 2010; Dessaud et al., 2007; Sasai et al., 2014).

The explants were treated with four different concentrations of Shh (0, 0.25, 1, and 4 nM), and the expression of transcription factors characteristic of dorsal-ventral domains was examined every 12 hours using immunofluorescence.

As a result, PAX7 expression, which marks the dorsal precursor domains, was detected in explants cultured without Shh or with 0.25 nM Shh at all time points. However, PAX7 expression was barely detectable at 1 nM and 4 nM of Shh (Fig. 1A). In contrast, the expression of OLIG2 and NKX2-2, which are expressed in two ventral precursor regions of pMN and p3, was induced by 1 nM and 4 nM Shh, respectively, at 12 hours (Fig. 1B,C). The expression of these factors was also detectable at 24 hours but was downregulated by 36 hours, suggesting a temporal presence of these neural progenitors.

**Fig. 1.**
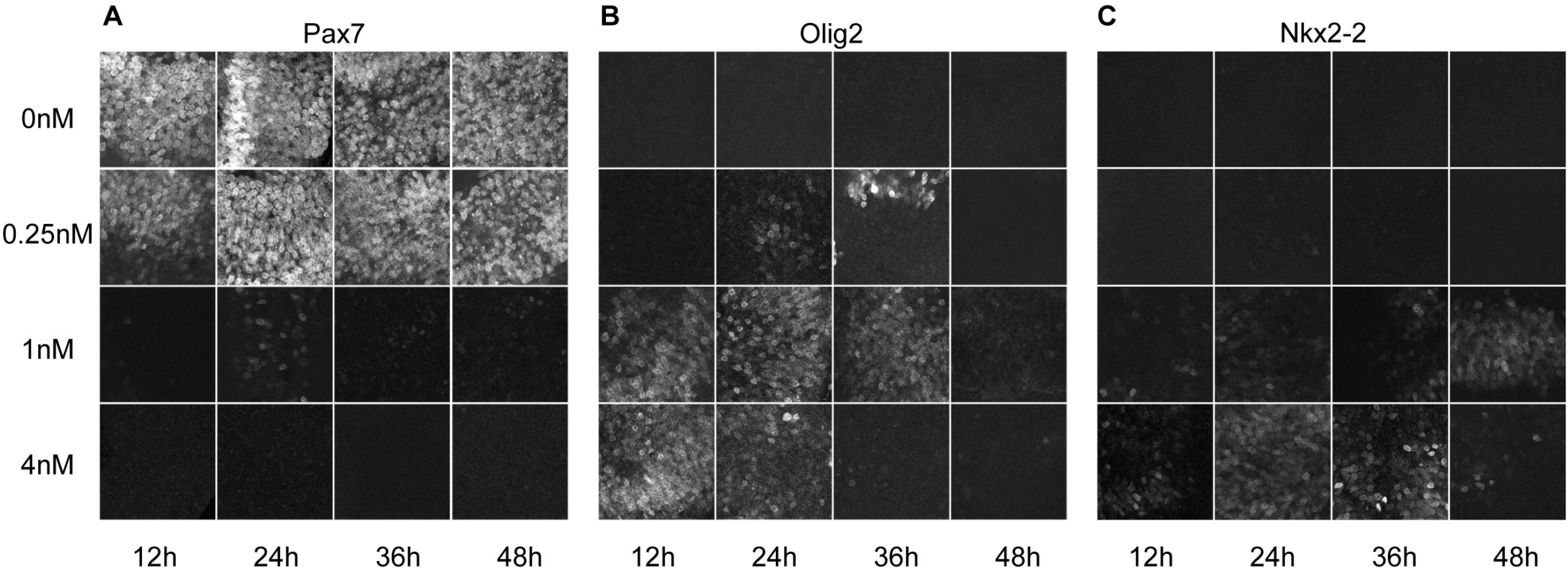
Differential expression of proteins along the D-V axis. Immunofluorescence staining of PAX7 (A), OLIG2 (B) and NKX2-2 (C) under the culture conditions of 0, 0.25, 1 and 4 nM of Shh for 12, 24, 36 and 48 hours.

Together, neural explants can recapitulate different identities and differentiation stages depending on the concentration and duration of Shh exposure.

### High-throughput expression profiling recapitulates the expression of genes induced by Shh

We next aimed to characterize the genome-wide expression profiles associated with Shh-induced neural progenitor differentiation. To achieve this, we extracted RNA from neural explants treated under 16 different conditions with different Shh concentrations and culturingperiods, as well as from explants immediately after preparation, which served as a control. The RNA samples were analyzed using high-throughput mRNA sequencing to profile gene expression. We represented the expression profiles of key transcription factors involved in neural differentiation and patterning as a three-dimensional graph illustrating the relationship between concentration, time, and expression levels (transcripts per million; TPM).

ARX (Aristaless related homeobox) has been shown to be expressed in the floor plate, with its expression being induced at 48 hours when explants were cultured with 4nM Shh (Sasai et al., 2014) (Fig. 2A). This observation aligns with previous findings where ARX was similarly induced under these conditions (Ribes et al., 2010; Sasai et al., 2014). FOXA2, which is also expressed in the floor plate and acts as an upstream factor for ARX (Sasai et al., 2014), contains Gli-binding sites (Peterson et al., 2012; Sasaki et al., 1999). Specifically, FOXA2, but not ARX, is directly induced by Shh. FOXA2 expression was observed as early as the 12-hour culture condition (Fig. 2B) and persisted, subsequently inducing ARX expression (Ribes et al., 2010; Sasai et al., 2014) (Fig. 2B).

**Fig. 2.**
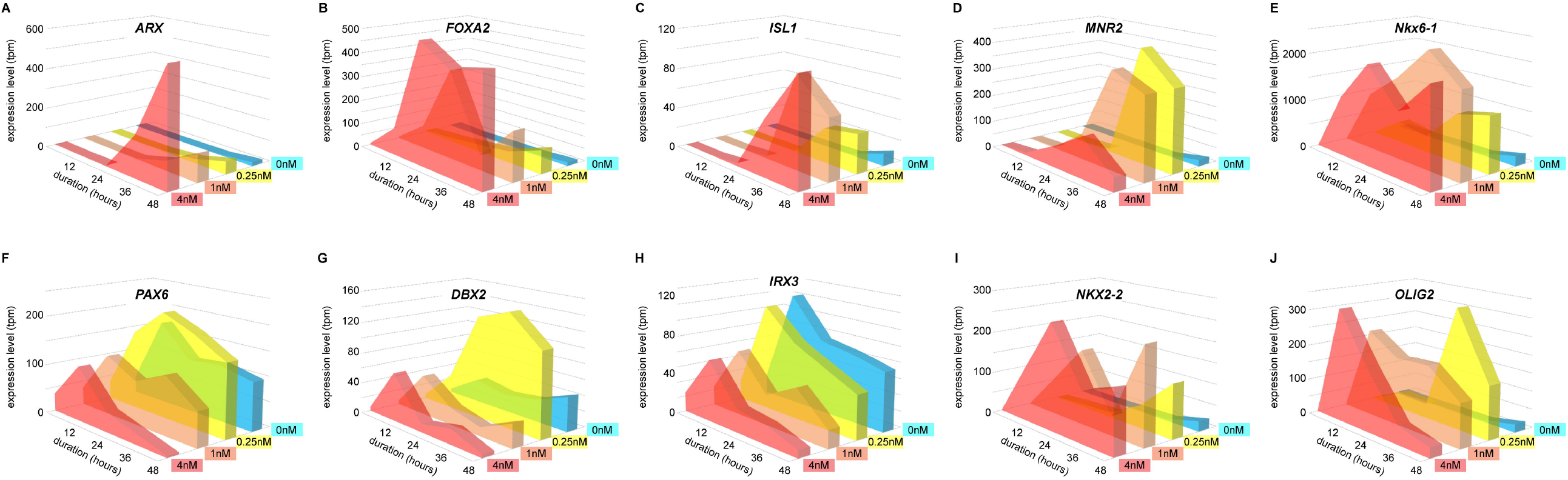
Three-dimensional graph illustrating the gene expression levels with different Shh concentrations and time points. The gene expression of *ARX* (A), *FOXA2* (B), *ISL1* (C), *MNR2* (D), *NKX6-1* (E), *PAX6* (F), *DBX2* (G), *IRX3* (H), *NKX2-2* (I) and *OLIG2* (J) is shown under culture conditions of 0, 0.25, 1 and 4 nM of Shh for 12, 24, 36 and 48 hours.

ISL1 (Islet-1), which is expressed in the motor neuron (MN) domain, one of the ventral neuronal regions (Pfaff et al., 1996), began to be expressed at 36 hours in response to 1nM and 4nM Shh, whereas 0.25nM Shh had little effect (Fig. 2C). MNR2 (HB9), an upstream regulator of ISL1 (William et al., 2003), exhibited a similar expression pattern (Fig. 2D). NKX6-1, which is widely expressed in the ventral neural domains (p2-FP progenitor domains), also showed early gene induction by Shh (Sander et al., 2000) (Fig. 2E). In contrast to the Shh-induced transcription factors, the expression of PAX6, which is expressed in the intermediate neural domains (dp1-pMN progenitor domains with varying expression levels), was repressed by Shh (Fig. 5F); therefore, its expression was significantly downregulated as the Shh concentration increased (Fig. 5F), despite a slight increase at the 12-hour time point. Additionally, it is noteworthy that DBX2 (Developing Brain Homeobox 2), expressed in the dp5-p1 progenitor domains, was primarily observed only under the 0.25nM condition, which is consistent with the functional analysis where DBX2 has a mutually repressive relationship with NKX6-1 (Briscoe et al., 2000) (Fig. 2G).

IRX3 has been shown to be expressed in the dp1-p2 progenitor domains (Briscoe et al., 1999), with higher levels of expression observed at 0nM and 0.25nM, and a transient upregulation at the 12-hour time point (Fig. 2H).

The two transcription factors, NKX2-2 and OLIG2, expressed in the p3 and pMN progenitor domains, respectively, have GBSs (Oosterveen et al., 2012; Peterson et al., 2012), and are directly upregulated by Shh. Although both genes were upregulated at 12 hours at 1nM and 4nM, OLIG2 expression quickly decreased at 24 hours at 4nM, presumably due to repression by NKX2-2 (Balaskas et al., 2012) (Fig. 2I,J), which leads to the formation of the p3 progenitor domain (Dessaud et al., 2007). Overall, the dataset describing the genome-wide expression profiles aligns with in vivo expression patterns and systematically demonstrates the morphogen model, where the acquisition of different neuronal characteristics depends on signal concentrations.

### Genes were categorised into 224 patterns according to their expression profiles

We aimed to discover novel GRNs by applying known GRNs to the gene expression dataset described above. To achieve this, we first compiled GRNs that had been identified through traditional experimental methods, including chromatin immunoprecipitation (ChIP), overexpression, and loss-of-function experiments (Sagner and Briscoe, 2017). Fig. 3A illustrates these GRNs, where green arrows represent induction relationships and red stop arrows denote inhibition relationships. This network is further visualized in matrix form in Fig. 3B, where the genes arranged vertically are regulated by those arranged horizontally. In subsequent analyses, we will use this matrix representation to elucidate the dependency relationships between genes.

**Fig. 3.**
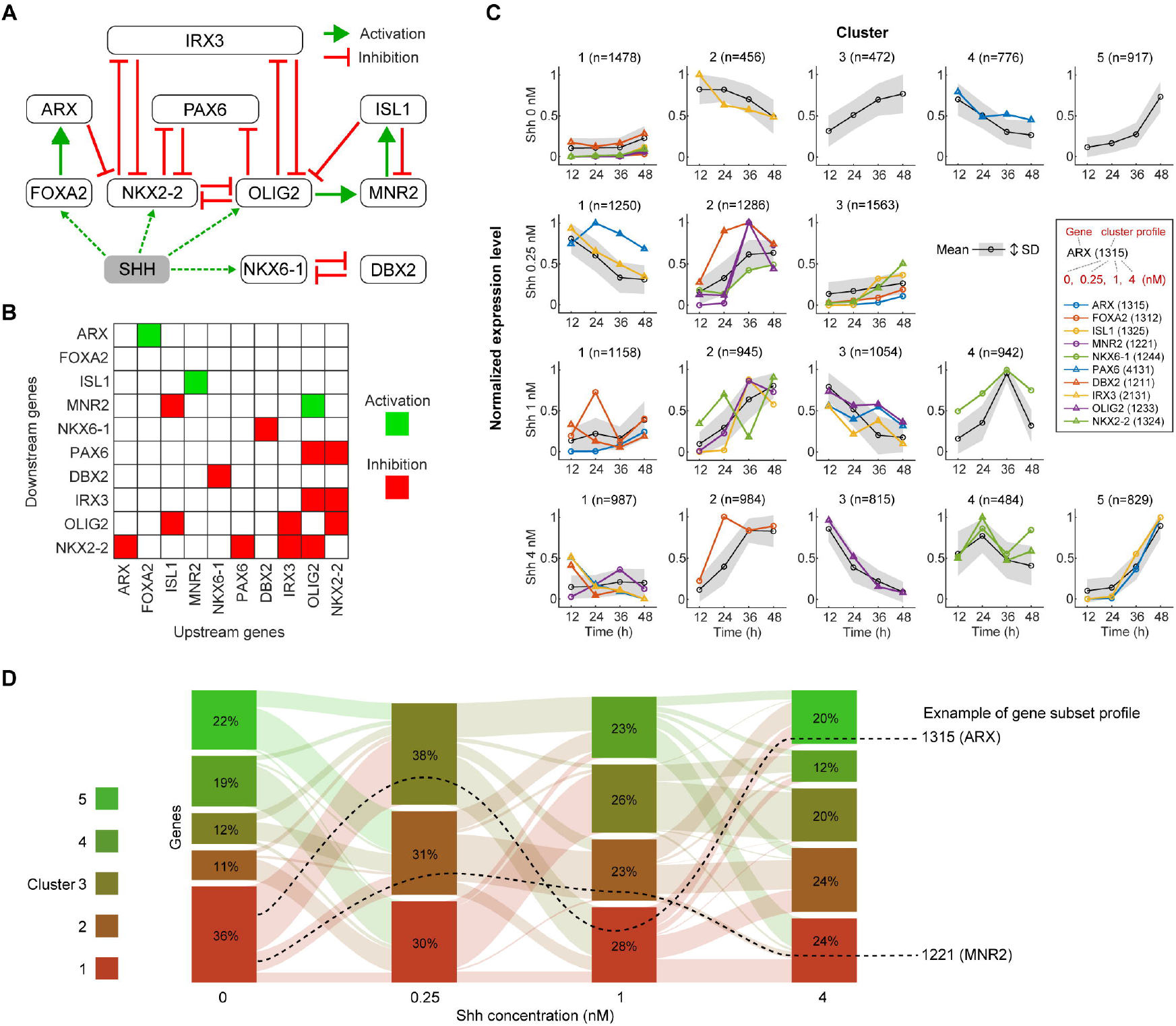
Known genes, pathways and clustering of expression patterns. (A) Diagram of the ten known genes involved in neural tube formation and the established regulatory pathways between them. (B) Matrix representation of the pathways shown in (A), with downstream genes arranged vertically and upstream genes arranged horizontally. (C) Shh concentration-dependent clustering of the expression time series for the 4,103 genes after preprocessing. The bottom row represents the highest Shh concentration, corresponding to the ventral side of the neural tube. The time series of the 10 known genes from Fig. 2 are included within one of these clusters. Each gene is assigned a cluster number for each concentration, resulting in a four-digit cluster profile for each gene (Fig. S1D). (D) Sankey plot showing the proportion and flow of genes belonging to each cluster in each concentration condition. Each gene is profiled using the cluster number at the four concentrations. As an example, the flows of ARX and HB9 are shown with a black dotted line.

First, we extracted genes that exhibited significant expression based on three criteria. We initially filtered the genes by requiring that the maximum expression level across the 16 conditions was significantly high and showed significant variation over time (Fig. S1A). We further refined the selection to include only those genes with an absolute log2 fold change of at least 1 compared to the control (Fig. S1B). This process resulted in the selection of 4,103 genes for further analysis.

Next, we classified the genes based on their expression time series. Genes with similar expression patterns are challenging to distinguish, so they were grouped together. For each Shh concentration condition, we applied k-means clustering to the time-series data of four time points. The optimal number of clusters was determined using the gap statistic (Fig. S1C). Consequently, each gene was classified into different clusters depending on the concentration (Fig. 3C). Based on these clustering results, we further subdivided the gene groups. Each gene was assigned a profile number corresponding to its cluster assignment across the four concentrations, forming a four-digit label. For example, ARX was assigned the profile 1315, while MNR2 was assigned the profile 1221 (Fig. 3D). This profiling produced a total of 224 gene subsets (GSs; Fig. S1D). These findings demonstrate that genes belonging to the same cluster at one concentration may belong to different clusters at other concentrations, indicating that although the GRN remains the same, its functional dependencies vary with Shh concentration. In this study, we used the average time series of these GSs (Fig. 3C) to infer unknown genes and pathways involved in neural tube formation.

### Pathway estimation using the Bayesian framework and validity evaluation of candidate genes

Since there are a certain number of genes and pathways with known involvement, in this study, we considered these known genes and pathways as a core network, and estimated the genes that are required but whose functions are currently unknown (unknown genes). Specifically, we developed a Bayesian framework called network assembly and inference using stepwise testing with optimization (NAISTO) that incorporates unknown genes one by one into the core network and performs pathway estimation and validity evaluation.

The algorithm comprises two stages: pathway estimation and the evaluation of the validity of added unknown genes. First, a linear regression model was constructed based on GSs containing 10 known genes. During this process, one of 214 unknown GSs was selected and added to the regression model. The regression equation included both known and unknown pathway parameters, with prior constraints of non-zero values and sparsity applied to each parameter group (Fig. 4A). We then calculated the posterior distribution using the model’s likelihood and the pathways’ priors, and estimated the parameters that maximized the posterior distribution’s probability density. Subsequently, we utilized the matrix representation of the network to check the presence of any known pathways. This process was repeated for each of the 214 unknown GSs (Fig. 4A). The validity of the unknown GSs was evaluated using the estimated pathway parameters (Fig. 4B). The evaluation criteria were based on two conditions: the conservation of known pathways and the sparcity of unknown pathways. In the ranking coordinate system, the closer the GS is to the origin (i.e., the smaller the *F*_*i*_), the more likely it is that the GS contains a regulatory gene (Fig. 4C).

**Fig. 4.**
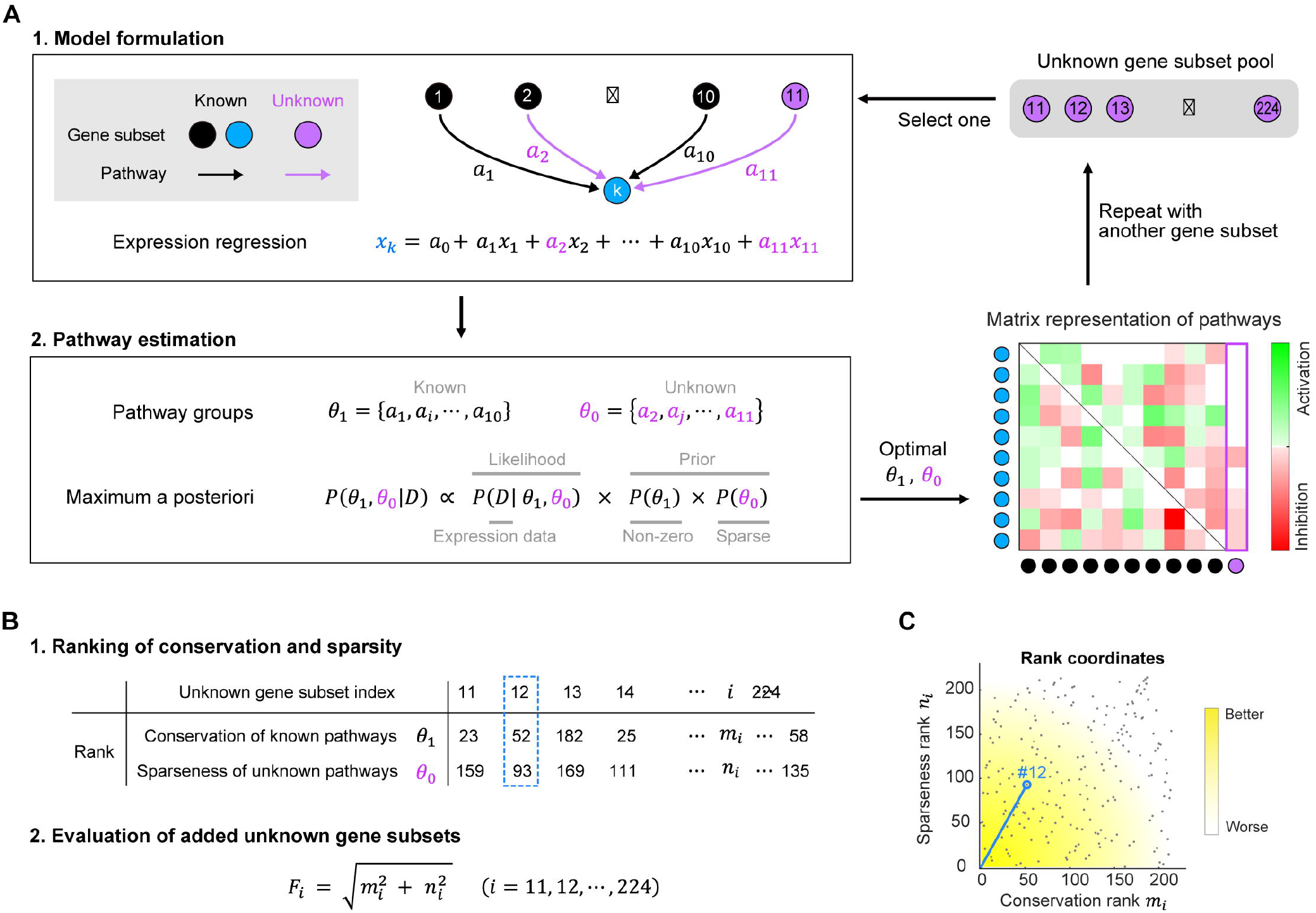
Algorithm for estimating genes and pathways of unknown involvement in neural tube development. (A) Formulation and pathway estimation process involves adding and replacing one unknown GS to the network of known GSs. Each known GS is modeled using linear regression, with 9 known GSs and 1 unknown GS to describe the relationships among the known GSs, including the unknown pathway. The parameters of the regression equation reflect the presence or absence, activity, and inhibition of pathways, and these are categorized into known (*θ*_*k*_) and unknown (*θ*_0_) in the pathway estimation. Non-zero prior knowledge is applied to *θ*_*k*_ and a sparse prior to *θ*_0_, and the parameters that maximize the posterior distribution are estimated using Bayes’ theorem. To verify the impact of prior information, the estimation results are visualized in the matrix representation as shown in Fig. 3B. This process is repeated for each unknown GS selected from the pool. (B) An evaluation metric for the validity of the unknown GS added in (A). For the pathways estimated using each unknown GS, a higher median value of the known pathway parameters indicates better conservation, while a greater number of small unknown pathway parameters (0.05 or less; see Methods) indicates higher sparsity. Each unknown GS (from No. 11 to No. 224) is ranked according to these metrics, and the square root of the sum of the squares of the ranks (*F*_*i*_) is used as a validity evaluation. (C) Coordinate representation of the ranking. The location of the unknown GS is plotted. The closer a GS is to the origin (indicated by yellow), the more valid it is. For example, the 12th unknown GS, marked by a blue rectangle in the table, corresponds to the blue circle in the rank coordinates, and its distance from the origin is *F*_12_.

**Fig. 5.**
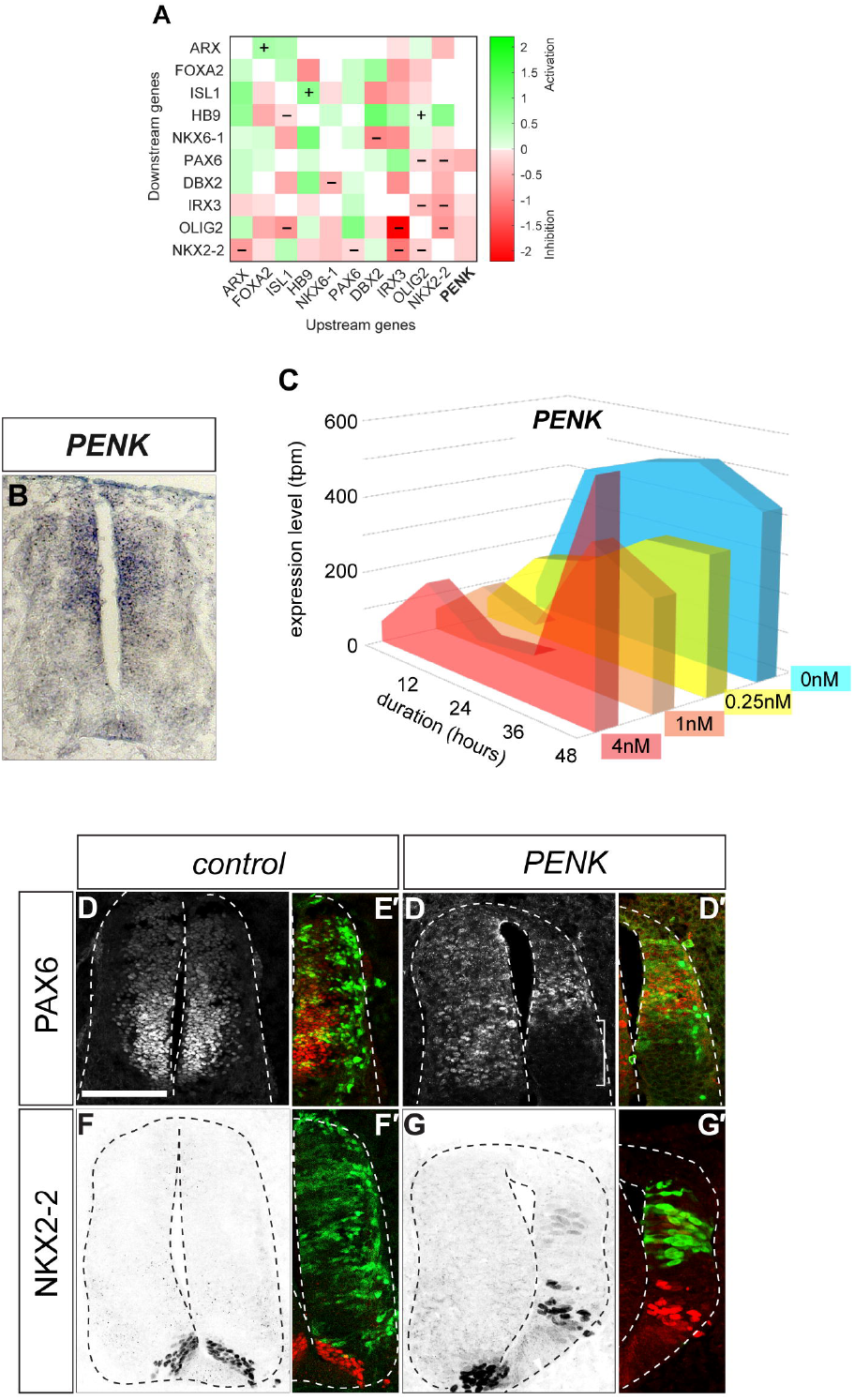
Validation of one of key candidate genes identified through the estimation. (A) A matrix where PENK is involved in the GRNs, with PENK assumed to repress the genes PAX6, IRX3, OLIG2, and NKX2-2. (B) *PENK* is highly expressed in the dorsal region of the neural tube, as detected by in situ hybridization. (C) The mRNA levels of *PENK* at each condition (Fig. S1). (D-I’) The effect of *PENK* overexpression. Expression plasmids for the *control* (D,D’,F,F’) or *PENK* (E,E’,G,G’) were electroporated at HH stage 12, and the expression of PAX6 (D-E’) and NKX2-2 (F-G’) was detected by immunofluorescence at 48 hpt. Scale bar in (D) = 100 μm, which applies to (D-G’).

To determine the appropriate degree to which the non-zero and sparse prior constraints should be applied, it is necessary to adjust the weight coefficients. To achieve this, we performed leave-one-out cross-validation on the 10 known GSs (see “Optimization of the weights of the objective function” in Methods). In this process, one of the 10 known GSs was left out and considered as unknown, mixed with the 214 unknown GSs, and the weight coefficients were adjusted so that the sum of the evaluation index *F*_*i*_ of the left-out GS would be minimized. Specifically, a grid search was conducted based on the distribution shape of non-zero pathways (Fig. S2A) and the coefficients of known pathway conservation and unknown pathway sparsity (Fig. S2B). With the optimal weight coefficients, all 10 known GSs were ranked highly (Fig. S2C). The regression time series and parameter matrix obtained from these weight coefficients are summarized in Fig. S3. These results show that this method appropriately ranks important GS at the top of the list.

In this way, we ranked the 214 unknown GSs using the optimized weight coefficients to estimate the 11th regulatory factor. The top 10 GSs, along with the gene names contained in each GS, are shown in Tab. 1.

**Table 1.** Top 10 GSs extrapolated from the estimation.

| Rank | 1 | 2 | 3 | 4 | 5 | 6 | 7 | 8 | 9 | 10 |
| --- | --- | --- | --- | --- | --- | --- | --- | --- | --- | --- |
| Genes | SLC39A13 | SDK2 | LOC107049387 | DNAJC5G | **PENK** | CASP2 | C14H16orf45 | EPN3 | LOC112532802 | BRWD3 |
|  | FKBP10 | ARHGAP39 | LOC107049379 | SAYSD1 | LOC107049496 | KIAA1217 | DPYSL3 | FAT3 | GDF3 | STRN |
|  | PDIA6 |  | FHOD3 | TBC1D19 |  | LOC107056693 |  | TEAD1 | ACAT1 | COG1 |
|  |  |  |  | GATD3A |  | EFNA2 |  | EPCAM |  | SNRK |
|  |  |  |  | ERP29 |  |  |  |  |  | NCEH1 |

### Validating the network through overexpression analysis in chick embryos

We sought to validate the gene functions predicted by our computational analysis. Among the highly ranked candidate genes listed in Tab. 1, we specifically searched for potential signaling factors and transcription factors, as these are likely to influence the expression of other genes. We chose to focus on the gene *Proenkephalin* (*PENK*) because it encodes a secreted peptide.

First, we predicted the activity of PENK. By incorporating PENK into the gene network, we predicted the expression of various genes through a linear combination. The results indicated that, in addition to the expression of PAX6 and IRX3, the expression of OLIG2 and NKX2-2 would also be suppressed (Fig. 5A).

To validate this prediction, we examined PENK expression in the neural tube at HH stage 22, corresponding to 36-48 hours of development in explants. As predicted by the gene expression data, PENK expression was observed in the dorsal domains of the neural tube (Fig. 5B). This expression pattern is consistent with the trend in the database, where PENK is highly expressed in regions with low or absent Shh levels (Fig. 5C).

Next, we electroporated an expression vector containing the coding region of PENK into the neural tube of chick embryos at HH stage 12. The embryos were harvested 48 hours post-electroporation, and the expression of PAX6 (Fig. 5D-E’) and NKX2-2 (Fig. 5F-G’) was analyzed. The results showed a downregulation of PAX6, while NKX2-2 were ectopically induced. These observations partially consistent with our predictions, as PAX6 expression was indeed downregulated. However, the upregulation of NKX2-2 was unexpected (Fig. 5F-G’). We hypothesized that this discrepancy might be due to the artifact effect of the PENK overexpression, which could have aberrantly suppressed PAX6.

Thus, one of the genes found from the gene regression data was found to be certainly active and is involved in the D-V patterning of the neural tube.

## Discussion

### Shh-treated neural progenitor cells are useful for identifying the causal relationship between signal inputs and gene expression

In this study, we successfully recapitulated positional information in neural progenitor cells by applying varying concentrations of signaling factors and adjusting culture durations, thereby inducing the identity of the ventral neural tube. While previous single-cell expression analyses have elucidated gene expression profiles associated with each neural progenitor domain during neurogenesis, they have been limited in directly linking the input from signaling factors to gene expression output (Delile et al., 2019). Our explant-based gene expression analysis offers a significant advantage by providing two-dimensional data in terms of Shh concentration and differentiation time, enabling a direct correlation between input and output. In addition, the ability to subdivide expression patterns (gene subsets) based on Shh concentration provides significant insights for the estimation of unknown genes and pathways. This approach can be applied in future studies to track gene networks involved in the differentiation of tissues other than neural tissue and to discover novel signals for differentiation.

To understand the importance of specific genes in organ differentiation, common methods involve forced expression or knockout of the gene of interest (Balaskas et al., 2012). While this approach is effective in experimentally demonstrating gene significance based on phenotypic outcomes, it carries the risk of phenotypic variation depending on the experimental system (e.g.; model animals or experimental methods), which could lead to misinterpretation due to artifacts. In contrast, in silico network prediction allows for the flexible manipulation of gene expression levels, facilitating the prediction of phenotypic outcomes. The integration of known networks with extensive expression data in the future is expected to enable more precise predictions of gene functions.

The fact that neural progenitor cells exhibit different responses to varying Shh concentrations suggests the presence of sophisticated information processing mechanisms within the cells. Particularly, it is intriguing to note that each gene’s influence on others (either activation or repression) is consistent, irrespective of Shh concentrations. This ability that the same network elicits different responses to different stimuli is reminiscent of how neural circuits generate distinct outputs based on specific inputs. In neuroscience, such dynamic networks are referred to as “functional connectivity” and are one of major focuses of current research (Deco et al., 2014; Honey et al., 2009; Hutchison et al., 2013). A single input stimulus is insufficient to elucidate dynamic systems. Therefore, it is appropriate that this study obtained expression data at multiple Shh concentrations.

### Perspectives for the enhancement of gene prediction methods based on known genes and pathways

The approach of adding one unknown gene at a time and evaluating its validity proved to be effective in this study. A key factor contributing to this success was the availability of substantial information on known genes and pathways. If the known information had been limited, the estimations would have faced challenges due to the increased degree of freedom. However, to further enhance the effectiveness of this method, it is essential to prepare a more detailed dataset. For instance, increasing the number of Shh concentration conditions and further subdividing the gene subsets could improve the accuracy of predictions. While this study successfully identified the 11th gene, it is anticipated that the 12th and subsequent genes could be discovered using the same step-by-step approach. This would not only expand the existing knowledge base but also likely lead to further improvements in prediction accuracy. The method employed in this study is based on Bayesian statistical learning and optimizes the weights for pathway constraints. However, several options exist for selecting candidate genes, which may vary depending on the dataset. For instance, a more effective indicator might be introduced for the ranking step used to evaluate the validity of added unknown genes. In this study, the median of the estimated parameters was used to assess the conservation of known pathways (Fig. 3B), but alternative statistics, such as the mean, could potentially be more effective depending on the specific data. The threshold value for considering the coefficients of unknown pathways to be zero (0.05 in this study) is also an adjustable parameter. These options can be validated through leave-one-out cross-validation.

### PENK is a novel component to regulate neural differentiation

In this study, we identified PENK as a novel gene involved in neural patterning and differentiation. PENK is expressed in the dorsal region of the neural tube (Fig. 5B). It encodes a precursor protein that is processed into several active peptides, including enkephalins, which play crucial roles in various physiological processes, such as pain modulation (Liu et al., 2023), stress response (You et al., 2023), immune function (Mas-Orea et al., 2023), neurotransmission (Wong et al., 2021), cardiovascular regulation (Emmens et al., 2021), endocrine function (Denning et al., 2008), and digestive system regulation (Kumar et al., 2024). However, the functions of PENK during embryogenesis remain unclear.

Enkephalins bind to opioid receptors (Tache et al., 2024) and activate several intracellular signaling pathways, including the inhibition of adenylate cyclase, the mitogen-activated protein kinase (MAPK) pathway, and the phosphoinositide 3-kinase (PI3K)/Akt pathway (Sobczak et al., 2014). Notably, enkephalins inhibit adenylate cyclase, leading to a reduction in intracellular cAMP levels. Since lower cAMP levels promote the ventralization of neural tube identity (Yatsuzuka et al., 2019), this mechanism likely reduces PAX6 expression while expanding ventral neural identities (Moore et al., 2016; Yatsuzuka et al., 2019). However, given that PENK is expressed in the dorsal region of the neural tube, the mechanisms by which it contributes to neural patterning are puzzling. Further in-depth analyses, such as loss-of-function experiments, are needed to uncover the essential roles of PENK in neural development.

## Methods

### Ethical statement

All animal experiments were performed under the approval of the Animal Welfare and Ethical Review Panel of Nara Institute of Science and Technology (approval numbers of 1810 and 2311) with the protocols in accordance with the national and internal regulations.

### Preparation of chick neural explants and high-throughput expression profiling

Chicken eggs used in this study were purchased from Yamagishi Farm (Wakayama Prefecture, Japan) and incubated in a humidified incubator at 38°C. Neural explants were isolated from the preneural tube area in the caudal region of HH stage 9 embryos (Delfino-Machin et al., 2005) and embedded in collagen gel (SIGMA; C4243) (Sasai et al., 2014). The explants were cultured in DMEM/F-12 medium (Wako; 048-29785) supplemented with mito-serum extender (BD; 355006) and a penicillin/streptomycin/glutamine mixture (Wako; 161-23201) for the specified time periods. The Shh protein was produced in house (Kutejova et al., 2016; Yatsuzuka et al., 2019). The concentration of 4 nM (highest concentration) was defined as the level that induces floor plate cells positive for ARX expression at 48 hours (Sasai et al., 2014). Explants cultured without Shh tend to acquire intermediate and dorsal neuronal characteristics.

### High-throughput expression analysis

Total RNA was extracted from 20-30 explants using a NucleoSpin RNA extraction kit (U955C, Takara). mRNA sequencing analysis was performed by BGI (Beijing, China). The sequencing platform was BGISEQ, and 100-base paired-end sequencing was performed. Approximately 20 million reads were obtained for each sample, and alignment was performed using Bowtie2. A total of 16,416 genes were aligned.

### Electroporation of PENK

The coding region of the PENK gene was subcloned into the pCIG vector. The empty or the plasmid carrying the PENK gene were electroporated at HH stage 12 and were harvested at 48 hours post transfection (hpt).

### Expression analysis

In situ hybridization and immunofluorescence were performed as previously described (Yatsuzuka et al., 2019).The antibodies used in this study were; PAX6 (1:1,000 rabbit; Millipore 0AB2237), PAX7 (1:50 mouse; DSHB 0PAX7), NKX2-2 (1:50 mouse; DSHB 074.5A5), NKX6.1 (1:50 mouse; DSHB 0F55A10), OLIG2 (1:1,000 rabbit; Millipore 0AB9610) and GFP (1:1,000 sheep; AbD Serotec 04745-1051). The secondary antibodies were; anti-sheep IgG (1:500 Jackson 0713-096-147), anti-mouse IgG (1:500 Jackson 0 715-166-151) and anti-rabbit IgG (1:500 Jackson 0711-166-152).

### Data pre-processing

#### Filtering of significantly expressed genes

First, we selected significantly expressed genes based on the distribution of the maximum read count (TPM) across these 16 conditions (Fig. S1A, left panel). Given that the distribution was bimodal, we approximated it using a mixture of exponential and normal distributions, selecting genes that exceeded the mean minus one standard deviation of the normal distribution. Next, to identify genes with significant variation across the 16 conditions, we calculated the coefficient of variation (standard deviation divided by the mean) for each gene and approximated its distribution by a kernel density estimation method. Genes with variation greater than the mode were selected (Fig. S1A, right panel). Finally, we selected genes with an absolute log2 fold change greater than 1 relative to the control (Fig. S1B). These filtering steps reduced the number of genes from 16,416 to 4,103.

#### Clustering and creation of Shh stimulus-dependent subsets

Since genes with similar expression time series were indistinguishable, we clustered the expression time series patterns across four time points for each Shh concentration. First, the expression levels at the 16 conditions were scaled for each gene to a minimum of 0 and a maximum of 1. Next, k-means clustering was performed on the four time points for each Shh concentration condition. The optimal number of clusters was determined by calculating the gap statistic for each condition (Fig. S1C). As a result, each gene was assigned a cluster number for each concentration, forming a unique four-digit integer profile (Fig. S2D). Based on these profiles, 224 gene subsets were created (Fig. S2E).

### Mathematical models and estimation algorithms

#### Regression equations for expression time series

We approximated the mean expression time series of the known GSs using linear regression, with the expression time series of other GSs as predictors. Using the known GSs and pathways as the core model, we included an additional unknown GS and pathway (model formulation in Fig. 4A). We defined the expression level of the *i*-th known GS at time *t* as *x*_*i*_(*t*) (*k* = 1, ⋯, 10; *t* = 1, ⋯, 16). When the expression level of the *k*-th GS was the objective variable, the nine known GS other than the *k*-th GS, along with the 11th additional unknown GS, were used as explanatory variables, as described by the following equation:

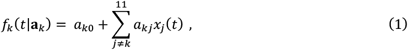

where the parameter vector *a*_*k*_ = [*a*_*k*0_, *a*_*k*1_, …, *a*_*k*11_] represents the intensity of the pathways and is estimated from the expression data. If any of these elements are estimated to be zero, it indicates the absence of corresponding pathway.

Neural tube progenitors functionally alter their regulatory networks depending on Shh concentration, but the regulatory relationships between genes remain consistent (Fig. 3A). In other words, each GS exhibits 16 distinct expression levels (across 4 concentration conditions and 4 time points), which are determined by a specific **a**_*k*_ independent of Shh concentration. Therefore, one specific set of **a**_*k*_ is estimated for all expression levels, regardless of concentration and time.

#### Pathway estimation using prior information

In this study, we divided all pathway parameters **a**_*k*_ (*k* = 1, …, 10) into known pathways (*θ*_1_) and unknown pathways (*θ*_0_) to impose constraints accordingly. We then estimated these pathways using a Bayesian statistical machine learning framework, assigning distinct prior probability distributions to the known and unknown pathways (see Fig. 4A for pathway estimation).

##### Likelihood of expression levels

Assuming that the error between the observed expression level *x*_*k*_(*t*) of the known GS at time *t* and the modeled expression level *f*_*k*_(*t*|**p**_*k*_) follows Gaussian noise, the likelihood function is given by:

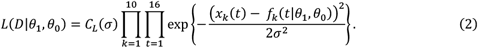

Here, *D* represents all observed expression levels {*x*_*k*_(*t*)}, *σ*^2^ is the variance of the noise, and *C*_*L*_(*σ*) is the product of the normalization constants for the Gaussian distribution, defined by *σ*.

##### *Prior* distribution *for known pathways*

To impose a non-zero constraint on the parameters *θ*_1_ of the known pathways, we modeled the absolute values of these parameters using a gamma distribution:

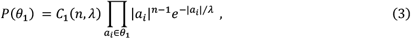

where *n* and *λ* are the shape and scale parameters, respectively, and *C*_1_(*n, λ*) is the product of the normalization constants for the gamma distribution, defined by *n* and *λ*. In the regression equation (Eq. 1), the sign of *a*_*i*_ is assigned according to the known activating or inhibitory effects.

##### *Prior* distribution *for unknown pathways*

Since only a few candidate unknown pathways are assumed to exist, we introduced a Laplace distribution as a sparse prior distribution for the parameters *θ*_0_:

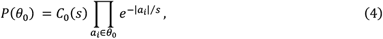

where *s* is the scale parameter, and *C*_0_(*s*) is the product of the normalization constants for the Laplace distribution, defined by *s*. This constraint has an effect equivalent to that of lasso regression.

##### Posterior distribution and objective function

According to Bayes’ theorem, the posterior distribution of *θ*_1_ and *θ*_0_ is given by:

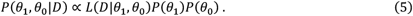

In this study, we sought the pathway parameters *θ*_1_ and *θ*_0_ that maximize the mode of this posterior distribution (maximum a posteriori). Substituting Eqs. 2-4 into the right-hand side of Eq. 5 and taking the logarithm, we obtain:

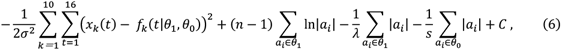

where the constant term *C* = ln *C*_*L*_(*σ*)*C*_1_(*n, λ*)*C*_0_(*s*) can be ignored because our interest lies in identifying the parameters that maximize the function, rather than in the maximum value itself. Multiplying all other terms by *α* = 2*σ*^2^ and flipping the sign, we derive the objective function to be minimized:

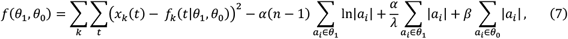

where *β* = 2*σ*^2^/*s*. The parameters that minimize this objective function:

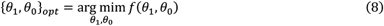

represent the parameter set {*θ*_1_, *θ*_0_} that maximizes the mode of the posterior distribution in Eq. 5. When a Gaussian distribution is used instead of a Gamma distribution, Eq. 7 becomes equivalent to the objective function of the elastic net algorithm.

The last three terms in Eq. 7 include the hyperparameters, *α, β, n*, and *λ*, which indicate their respective influence. A large *α* leads to stronger conservation of the known pathways, while a large *β* results in the estimation of as few unknown pathways as possible. However, if both *α* and *β* are too large, the first term in Eq. 7 is relatively undervalued, leading to larger errors between the observed and modeled expression levels. Therefore, it is necessary to optimize these four hyperparameters (see the section “Optimization of Objective Function Weights”).

#### Evaluation function for additional unknown GSs

We need an indicator to evaluate which unknown GSs should be added to the 10 known GSs. To compare the 214 unknown GSs, we assessed the conservation of the known pathway and the sparsity of the unknown pathway for the estimated pathway parameters when each unknown GS was added (ranking of conservation and sparsity in Fig. 4B).

We took the absolute value of the estimated parameter values for the known pathways, and used the median as the evaluation metric for pathway conservation. Based on this, we ranked the unknown GSs. The ranking for sparsity was based on the number of pathway parameter values less than or equal to 0.05. For the *i*-th candidate unknown GS, when the ranks for conservation and sparsity are *m*_*i*_ and *n*_*i*_ respectively, the evaluation value *F*_*i*_ of this GS is defined as:

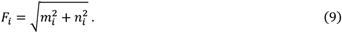

GSs with small *F*_*i*_ values are considered suitable as members of the GRN because they both conserve known pathways and introduce few additional unknown pathways (evaluation of added unknown GSs in Fig. 4B).

#### Optimization of the weights of the objective function

To optimize the hyperparameters (*α, β, n, λ* ; weight coefficients in Results section) in Eq. 7, we performed 10-fold cross-validation using 10 known GSs. In each iteration, we treated one known GS as unknown, mixed it with the 214 unknown GSs, and constructed a core network from the remaining 9 known GSs. For a given set of hyperparameters (*α, β, n, λ*), we formulated the objective function of Eq. 7 using the nine known and one unknown GS. This process was repeated 10 times, each time treating a different known GS as unknown. We then identified the combination of (*α, β, n, λ*) that minimized the sum of *F*_*i*_ across all 10 iterations, denoted by 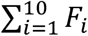. A grid search was conducted for the parameters (*α, β*) across four types of gamma distribution (Fig. S2A). As a result, the sum of *F*_*i*_ was minimized when the gamma distribution parameters were (*n, λ*) = (16, 0.1) and (*α, β*) = (0.04, 0.^2^) (Fig. S2B and C). These optimized hyperparameters were subsequently used to estimate the 11th unknown GS.

## Acknowledgements

The authors thank Jun-ichiro Yoshimoto, Toshiki Yoshimizu, Ryosuke Fujikawa, Tomohiro Kinugasa, and Yasumasa Bessho for comments, and the laboratory members for their discussions and supports.

## Author contributions

NS and YS conceived the study. NS prepared the neural explants and conducted the high-throughput expression analysis. YS devised the bioinformatics method, and CX and YS performed the analysis with assistance from TK and KK. NS conducted experiments to validate the predictions. YS, TK, KK, and NS secured funding. YS and NS wrote the manuscript.

## Availability of data and materials

The raw data for the RNA sequencing have been deposited at the DNA Data Bank of Japan (DDBJ) with the deposition number of PRJDB18433.

## Declarations

### Ethics approval and consent to participate

Not applicable.

### Consent for publication

Not applicable.

## Figure legends

**Supplementary Fig. S1 Preprocessing gene expression data and creating gene subsets**.

(A) Filtering genes by significance of expression level and variation. The distribution of the maximum expression level (TPM) across the 16 conditions was approximated by a mixture of an exponential distribution and a Gaussian distribution (red line in the left figure). We selected 9,293 genes using a threshold of 4.15, which is one standard deviation below the mean of the Gaussian distribution. Next, the distribution of the coefficient of variation—calculated as the standard deviation of the 16 values divided by the mean—was approximated using the kernel density estimation method (red line in the right-hand graph). We selected 7,162 genes using a threshold of 0.12, which corresponds to the mode of this distribution.

(B) Differential gene expression analysis against the control. Genes that exhibited more than a 2-fold increase or less than a 0.5-fold decrease in at least one of the 16 conditions were selected. A total of 4,103 genes were finally retained.

(C) Gap statistic analysis to determine the number of clusters in the expression time series for each concentration. The histogram shows the estimated number of clusters after 100 calculations.

(D) Histogram of the number of genes in each GS created in Fig. 3D, ordered by the number of genes. GSs that contain known genes are indicated by red bars, with the gene names written above the bars.

**Supplementary Fig. S2 Optimization of hyperparameters using leave-one-out cross-validation**.

(A) Four gamma distributions used as prior distributions for the known pathways. *n* and *λ* are the shape and scale parameters, respectively.

(B) Grid search for hyperparameters *α* and *β* when *n* = 16 and *λ* = 0.1. The z-axis represents the total sum of *F*_*i*_ values obtained from 10 cross-validation runs. The minimum value is indicated by a red dot.

(C) Sorted display of *F*_*i*_ values using the hyperparameters corresponding to the red dot in (B). The names of the known genes contained in the known GS that was left out in each cross-validation run are listed above the graph, and it was mixed with the other 214. The *F*_*i*_ of the left-out GS is indicated by a red circle in each graph. The number in parentheses next to each name represents the rank.

**Supplementary Fig. S3 Parameters estimated by cross-validation and comparison with time series**. In each section, the left-out known GS among the 10 is shown as a gray horizontal bar in the matrix representation. The left-out GS is also placed in the rightmost column of the matrix (blue rectangle) as an unknown upstream factor, and all pathways in the rectangle were estimated as unknowns under sparse constraints. The green and red colors in the matrix represent the estimated parameter values, with the plus and minus symbols indicating the known active and inhibitory pathways, respectively.

**Supplementary Tab. S1 The tpm representation of the gene expression of all genes in the neural explants cultured with the indicated conditions**.

**Supplementary Tab. S2 A graph visualizing gene expression**.

The expression trend of each gene can be found by copying and pasting each row from Tab. S1.

