## Supplementary figures and images for "Stepwise Bayesian Machine Learning Uncovers a Novel Gene Regulatory Network Component in Neural Tube Development"

### Supplementary Fig. S1

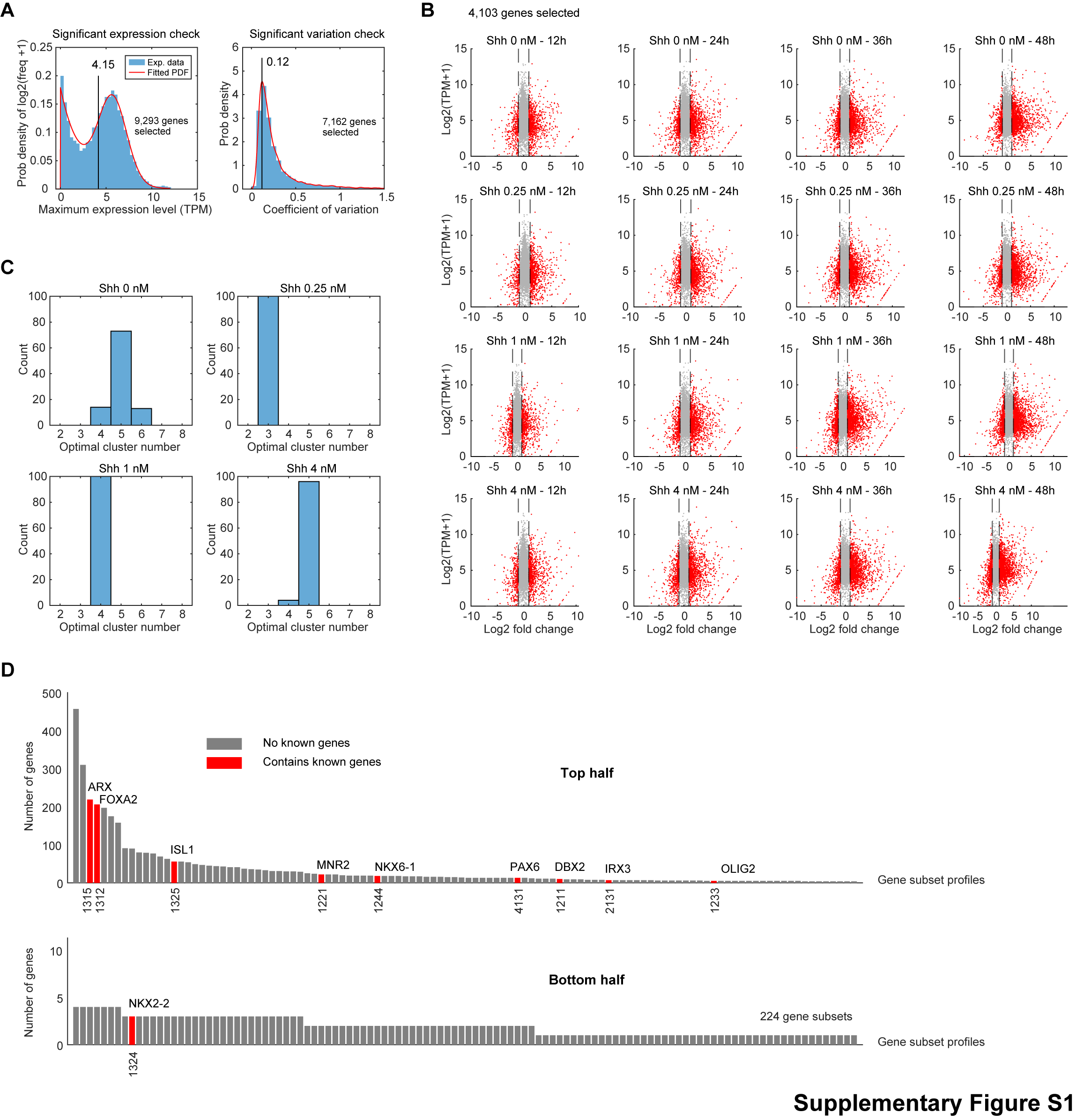

### Supplementary Fig. S2

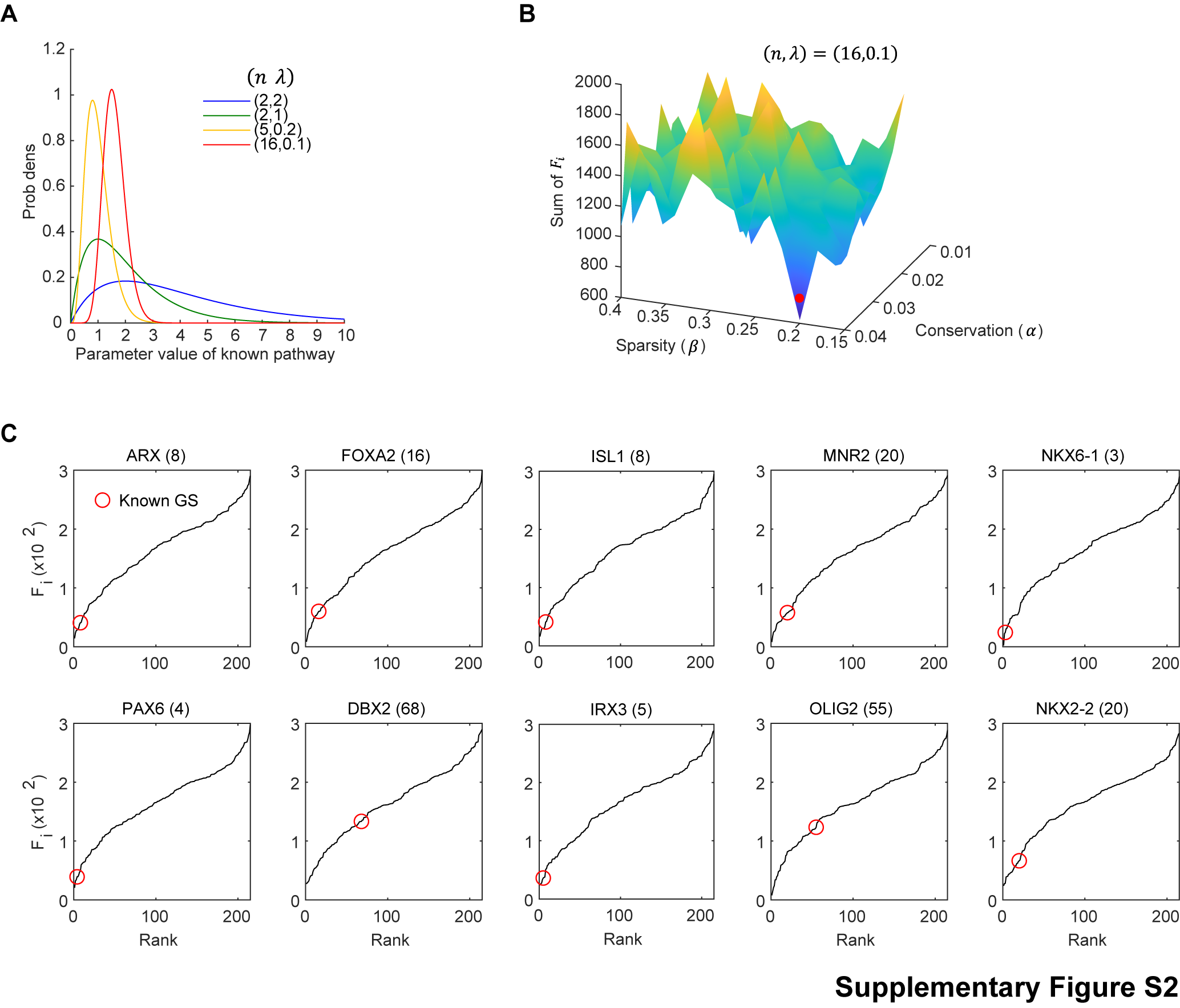

### Supplementary Fig. S3

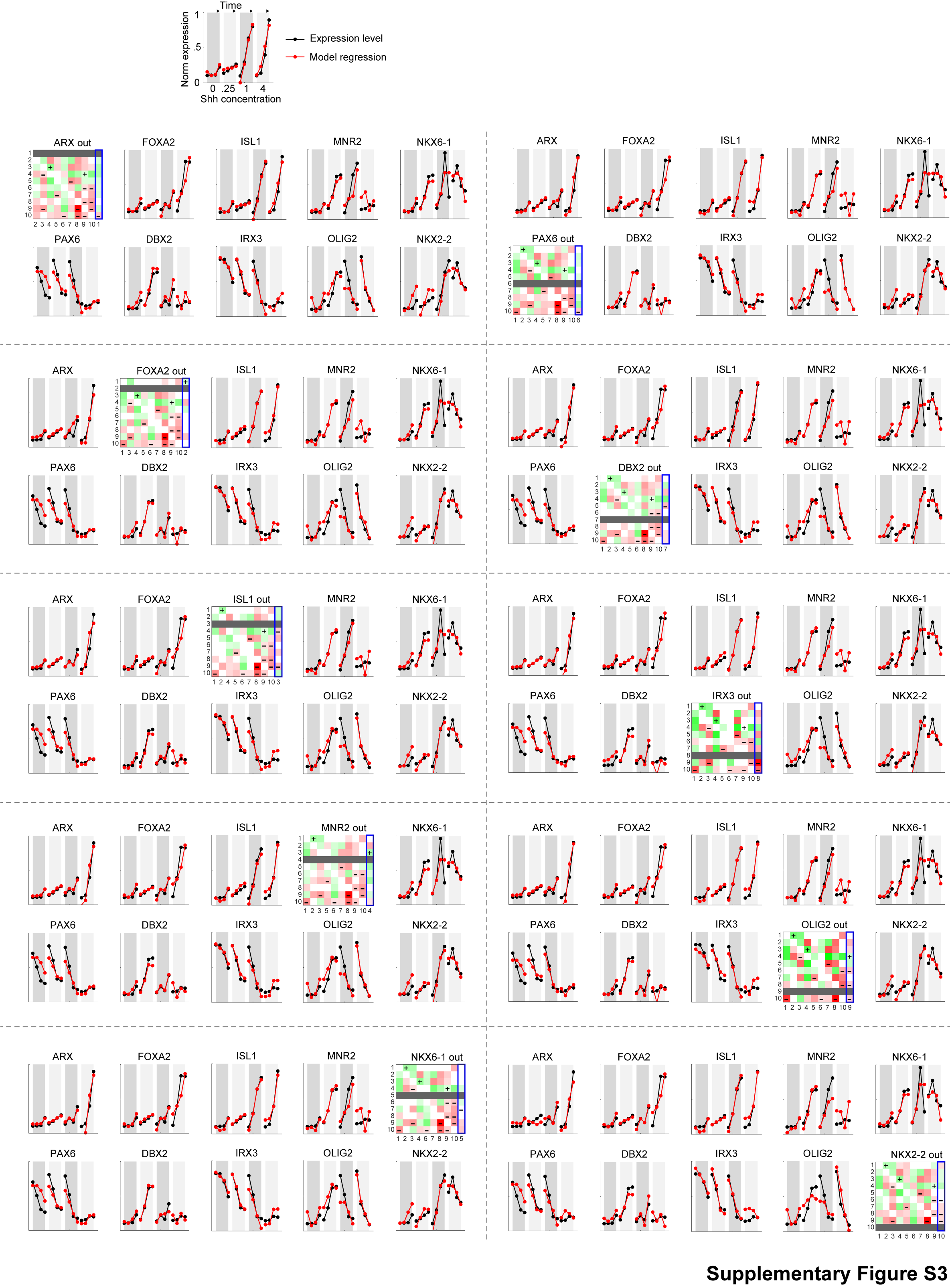
